# Site-specific machine learning predictive fertilization models for potato crops in Eastern Canada

**DOI:** 10.1101/2020.03.12.988626

**Authors:** Zonlehoua Coulibali, Athyna Nancy Cambouris, Serge-Étienne Parent

**Affiliations:** Department of Soils and Agrifood Engineering, Université Laval, Québec City, Quebec, Canada; Quebec Research and Development Centre, Agriculture and Agri-Food Canada, Québec City, Quebec, Canada

**Keywords:** Precision fertilization, Mitscherlich model, *k*-nearest neighbors, random forest, neuronal networks, Gaussian process, economic optimal dose, agronomic optimal dose, *Solanum tuberosum* L

## Abstract

Statistical modeling is commonly used to relate the performance of potato (*Solanum tuberosum* L.) to fertilizer requirements. Prescribing optimal nutrient doses is challenging because of the involvement of many variables including weather, soils, land management, genotypes, and severity of pests and diseases. Where sufficient data are available, machine learning algorithms can be used to predict crop performance. The objective of this study was to predict tuber yield and quality (size and specific gravity) as impacted by nitrogen, phosphorus and potassium fertilization as well as weather, soils and land management variables. We exploited a data set of 273 field experiments conducted from 1979 to 2017 in Quebec (Canada). We developed, evaluated and compared predictions from a hierarchical Mitscherlich model, *k*-nearest neighbors, random forest, neuronal networks and Gaussian processes. Machine learning models returned R^2^ values of 0.49–0.59 for tuber marketable yield prediction, which were higher than the Mitscherlich model R^2^ (0.37). The models were more likely to predict medium-size tubers (R^2^ = 0.60–0.69) and tuber specific gravity (R^2^ = 0.58–0.67) than large-size tubers (R^2^ = 0.55–0.64) and marketable yield. Response surfaces from the Mitscherlich model, neural networks and Gaussian processes returned smooth responses that agreed more with actual evidence than discontinuous curves derived from *k*-nearest neighbors and random forest models. When marginalized to obtain optimal dosages from dose-response surfaces given constant weather, soil and land management conditions, some disagreements occurred between models. Due to their built-in ability to develop recommendations within a probabilistic risk-assessment framework, Gaussian processes stood out as the most promising algorithm to support decisions that minimize economic or agronomic risks.

## 2 Introduction

Modeling provides a quantitative understanding of how crop systems operate [1]. Site-specific simulations of fertilizer requirements to obtain high local potato yield and quality rely on models’ ability to detect subtle variations in factors affecting plant growth and environment and to learn from the past to make predictions [2]. Several crop models have been developed with different degrees of sophistication, scale, and representativeness [2]. Mechanistic models have been published for potato cropping systems [3, 4]. Semi-mechanistic growth models could be used to downscale tuber yield assessment from regional to field levels [5, 6]. Multilevel modeling can assist in selecting a set of relevant parameters that impact tuber yield and fertilizer requirements, but can hardly predict site-specific nutrient requirements [7].

Several variables can impact fertilization at optimum tuber yield: soil type and quality [8, 9], organic fertilizers [10, 11], preceding crops [12–16], weather conditions [17], irrigation [18], timing, location and chemical form of the fertilizer applied [19], pests and diseases [20] and genetic factors such as cultivar longevity and growth rate [21, 22]. Air temperature, photoperiod, day length, intercepted radiation, water abundance, precipitations, root development and crop management were reported to be the driving variables for potato growth and development [8, 9, 23–26]. While the nitrogen (N) requirement of potato crops compares with other high N-demanding crops, phosphorus (P) uptake depends largely on close contact between roots and soil particles that, in turn, depends on soil texture, buffering capacity and moisture content [27, 28]. Due to a shallow system of fine roots and small biomass [29], especially in compacted soils [8, 9], potato is sensitive to nutrient and water stresses [30].

The N, P and K (potassium) requirements are thought to be cultivar- and market-specific [31–33]. Specific gravity (SG) is of particular concern for North-American processors [34, 35]. Other characteristics, such as tuber size and grade are also valued [34]. No model has yet addressed K requirements accounting for interactions between genetics, environment and management [36].

Growers tend to over-fertilize because of the potential economic loss from under-fertilizing [37, 38]. While N can cause nitrate contamination [39–42] and P the eutrophication of surface waters [43–45], K has no known deleterious effect on the quality of natural and drinking water. Attempts have been made to synthesize the results of fertilizer experiments using meta-analysis to derive N optima for specific soil texture and pH groups [46] or multilevel modeling combining soil, climate indices and management variables [7]. Even where field trials could identify nutrient optima [47], such optima cannot be generalized to conditions different from those of particular experiments [48, 49].

Although experimental data grow continuously in size and quality, it is still beyond researchers’ ability to integrate, analyze and make the best-informed decisions. Machine learning is an emerging technology that can aid in the discovery of rules and patterns in large sets of data [50]. The technology bypasses intermediate processes otherwise explicitly explained by a mechanistic modeling system and makes predictions directly based on input data [51]. Machine learning methods can combine fertilizer dosage, genetics, environmental and land management variables to predict tuber yield and quality. Classical models such as Mitscherlich are limited to plant-nutrient relationships [52].

We hypothesized that (1) genetics, environment and local land management practices are the main drivers of fertilizer requirements, (2) *k*-nearest neighbors, random forest, neural networks and Gaussian processes are more accurate in predicting marketable yield than classical Mitscherlich predictive models, and (3) the machine learning algorithms are equally able to predict economic optimal or agronomic optimal fertilizer doses. The objective of this study was to develop, evaluate and compare the performance of machine learning models in predicting N, P and K requirements for potato yield and quality.

## 3 Methodology

### 3.1 Data set and field experiments

The Quebec (Canada) potato data set comprised fertilizer trials conducted since 1979 between the US border (45^th^ parallel) and the Northern limit of cultivation (49^th^ parallel). Trials were arranged as randomized complete blocks or factorial designs. Seed spacing averaged 0.915 m between rows and 0.305 m on the row. Median plant density was 36,000 plants ha^-1^ in N trials, 33,100 plants ha^-1^ in P trials, 36,400 plants ha^-1^ in K trials, and 43,700 plants ha^-1^ in factorial NPK trials. No animal manure or compost had been applied in the spring and the preceding fall. The data set comprised 5,913 observations from the 208–273 field trials, depending on target variable (Table 1**Error! Reference source not found.**).

**Table 1:**
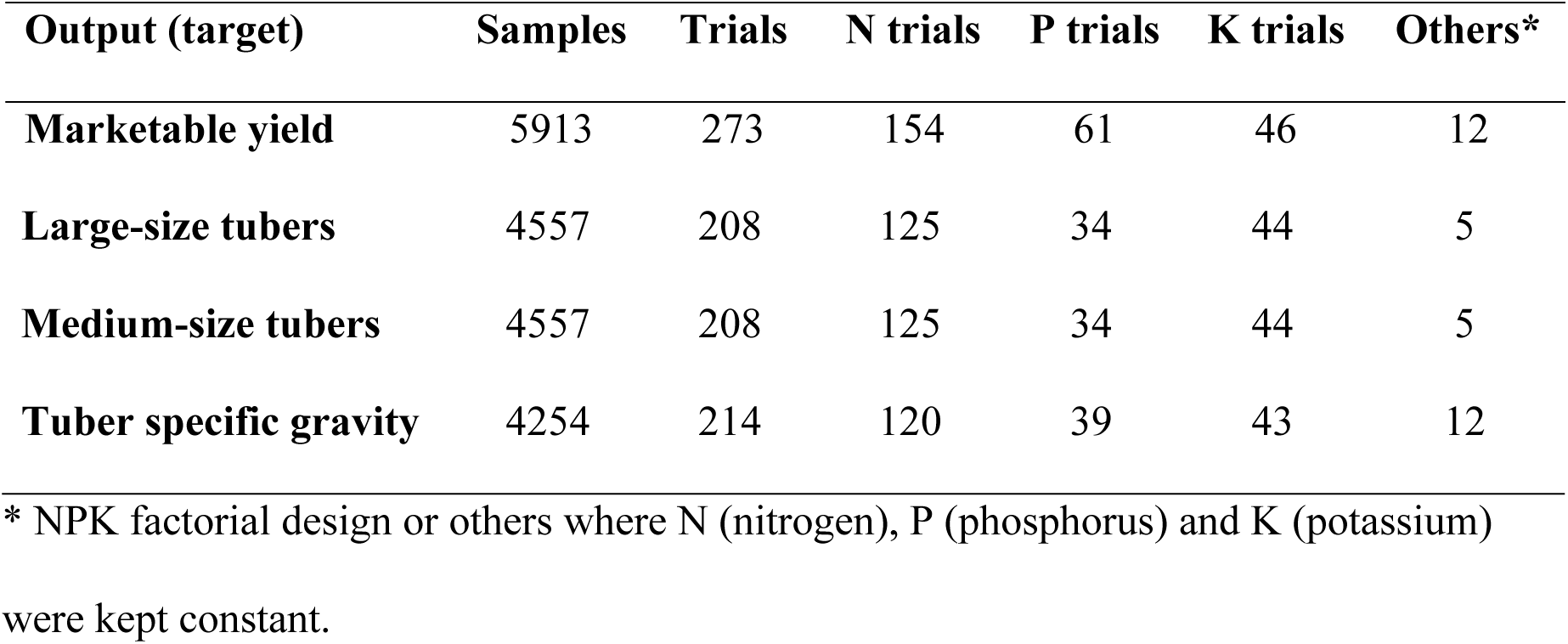
Description of the machine learning modeling data sets.

| Output (target) | Samples | Trials | N trials | P trials | K trials | Others* |
| --- | --- | --- | --- | --- | --- | --- |
| Marketable yield | 5913 | 273 | 154 | 61 | 46 | 12 |
| **Large-size tubers** | 4557 | 208 | 125 | 34 | 44 | 5 |
| **Medium-size tubers** | 4557 | 208 | 125 | 34 | 44 | 5 |
| **Tuber specific gravity** | 4254 | 214 | 120 | 39 | 43 | 12 |

\* NPK factorial design or others where N (nitrogen), P (phosphorus) and K (potassium) were kept constant.

There were 48 cultivars classified as early (65 – 70 days), early mid-season (70 – 90 days), mid-season (90 – 110 days), mid-season late (110 – 130) or late maturity (130 days and more) as suggested on the website of the Canadian Food Inspection Agency [53]. We matched the duration from planting to harvest but the classes names differed. The preceding crops were categorized as in Parent et al. [7] as grasslands, legumes, cereals, low-residue crops and high-residue crops. Toponymic names, geographical coordinates and years were recorded at each site. Fertilizers other than N, P or K, fertilizer source, dosage and application method, seeding density and date, harvest date, tuber marketable yield (excluding tubers < 2.5 cm in diameter), tuber size distribution (small, medium, large) and SG were recorded. The N fertilizers were either all applied at seeding or split-applied between seeding and hilling. The P fertilizers were banded at planting. The K fertilizers were band-applied or split-applied before planting and at planting. We added 17 trials conducted in 2016 and 2017 in the Outaouais, Centre-du-Québec, and Lac-Saint-Jean regions. We reported the growing season lengths provided by scouting teams covering the period from seeding to harvest and not strictly corresponding to the theoretical CFIA [53] growth duration as shown for cultivars Superior, Goldrush, Krantz and FL 1533 from the trials used for model analysis (Table 2).

**Table 2:** Cultivar and weather data from trials used for model analysis.

| **Trial number** | **194\*** | **8804** | **412** | **320** |
| --- | --- | --- | --- | --- |
| **Nutrient tested** | P | N | P | K |
| **Cultivar** | Superior | FL 1533 | Goldrush | Krantz |
| **Maturity class** | Early mid-season | Mid-season | Mid-season | Mid-season |
| **Growing season length (days)** | 102 | 131 | 108 | 112 |
| **Planting density (seeds ha<sup>-1</sup>)** | 36,430 | 43,716 | 36,433 | 31,226 |
| **Mean temperature (5 years) T°C** | 16 | 18 | 16 | 18 |
| **Total rainfall (5 years) mm** | 378 | 359 | 363 | 448 |

\* Trial n° 194 used for economic optimal and agronomic optimal doses computation. N: nitrogen, P: phosphorus, K: potassium

The experiments included four to six treatments arranged in a randomized complete block design with three replications of each treatment. Each experimental unit consisted of four or six rows measuring 6 or 8 m in length, with a row spacing of 0.915 m and within-row spacing varying with cultivar. The potato crop was planted in May (June in Outaouais), and tubers were harvested in September. The N doses varied from 0 to 260 kg N ha^-1^ with varying steps, and P was applied at a dosage of 0 to 130 kg P ha^-1^ with varying steps. K was applied at a dosage of 0 to 350 kg K ha^-1^ with varying steps. P and K fertilizers could be converted to P_2_O_5_ and K_2_O by multiplying P by 2.291 and K by 1.205. Other practices were managed uniformly by the grower.

At harvest, 3-m-long ridges in the middle two rows of each plot were dug and hand harvested. Tubers were divided into four categories as follows: culls, small (S), medium (M) or large (L), depending on the smallest diameter size measured with a ruler. The size cut-offs varied with cultivars and market. The marketable yield was calculated as total yield minus culls (tubers < 25 mm in size). Tubers with external defects such as secondary growth and soft rot were discarded. A representative sample of 20 medium-size tubers from each plot was used to determine tuber SG by the weight-in-air to weight-in-water method [54] as in equation 1:

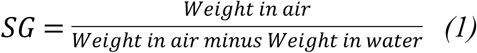

### 3.2 Soil profiles and analyses

The soils in our data set were classified according to the Canadian Soil Classification Working Group [55] and ordered along a gleyzation-podzolization gradient using tools of pedometrics [56]. Composite soil samples from the 0–20 cm layer were collected in the spring before planting to determine the initial soil physicochemical characteristics.

Grain size distribution was determined by sedimentation [57] or laser diffraction [58]. Where soil textural classes were not recorded, central values computed for sand, silt, and clay percentages (Table 3) using the Quebec soil data set [59] were assigned as proxies.

**Table 3.** Centroids of sand, silt, and clay percentages for soil textural classes derived from the Quebec soils data set [59].

| Textural class | Sand (%) | Silt (%) | Clay (%) |
| --- | --- | --- | --- |
| Coarse sand | 93.8 | 3.3 | 2.9 |
| Sand | 92.7 | 4.4 | 2.9 |
| Fine sand | 92.3 | 4.6 | 3.1 |
| Very fine sand | 92.9 | 4.4 | 2.7 |
| **Coarse loamy sand** | 82.5 | 11.3 | 6.2 |
| **Loamy sand** | 82.5 | 12.1 | 5.4 |
| **Fine loamy sand** | 81.4 | 14.4 | 4.2 |
| **Very fine loamy sand** | 80.7 | 11.2 | 8.1 |
| **Coarse sandy loam** | 66.2 | 20.4 | 13.4 |
| **Sandy loam** | 64.5 | 25.0 | 10.5 |
| **Fine sandy loam** | 60.4 | 31.6 | 8.0 |
| **Very fine sandy loam** | 60.2 | 32.8 | 7.0 |
| **Loam** | 42.4 | 41.1 | 16.5 |
| **Silty loam** | 18.2 | 67.0 | 14.8 |
| **Silt** | 6.0 | 88.1 | 5.9 |
| **Sandy clay loam** | 55.7 | 19.9 | 24.4 |
| **Clay loam** | 29.5 | 38.4 | 32.1 |
| **Silty clay loam** | 7.8 | 58.1 | 34.1 |
| **Sandy clay** | 50.2 | 11.2 | 38.6 |
| **Silty clay** | 2.7 | 48.1 | 49.2 |
| **Clay** | 11.9 | 34.9 | 53.2 |
| **Heavy clay** | 1.3 | 23.9 | 74.8 |

Soil pH was measured in water (1:1, v/v) or in a 0.01 M CaCl_2_ solution (1:1 v/v) [60]. Soil carbon concentration was determined using the Walkley-Black method [61] or Dumas combustion (Leco Instrument, Saint-Louis, MO). The two methods are closely related as in equation 2 [62]:

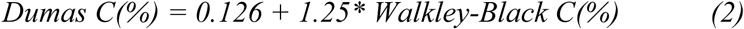

The pH_CaCl2_ was converted into pH_H2O_ where required, as in equation 3 [63]:

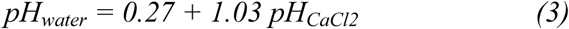

Soil P was extracted using the Mehlich-3 method [64] or Bray-2 converted to P Mehlich-3 values using the Khiari et al. [43] equation as in equation 4:

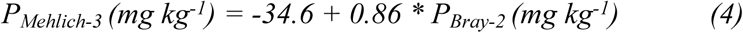

Soil Al was extracted using the Mehlich-3 method or, where not available, from the typical Al-Mehlich-3 value of soil series as reported by Tabi et al. [59]. Soil K, Ca and Mg were extracted using the ammonium acetate method or its closely-related Mehlich-3 extractant [65]. The P concentration was determined colorimetrically [66] or by inductively coupled plasma (ICP). The K concentration was determined by flame emission or ICP, and Ca, Mg, and Al concentrations were quantified by atomic absorption spectrometry or ICP.

The trials selected for model analysis (Table 4**Error! Reference source not found.**) showed soil pH levels ranging between the adequate limits of 5.2 to 6.2 for potato crops according to the *Centre de Référence en Agriculture et Agroalimentaire du Québec* [67]. The phosphorus saturation environmental index (P/Al)_Mehlich3_ classified the sites at extremely low environmental risk for P trials (1.4% to 1.6%), medium risk for N trial (11.1%) and very high risk for K trial (28.7%). Soil potassium levels showed extremely low (71.5 mg kg^-1^) and very low (83.1 mg kg^-1^) levels for P trials, medium level for K trial and high level for N trial [68].

**Table 4:** Median soil properties in selected fertilizer trials.

| Trial number | 194* | 8804 | 412 | 320 |
| --- | --- | --- | --- | --- |
| Tested nutrient | P | N | P | K |
| **Soil pH** | 5.5 | 5.5 | 5.8 | 6.1 |
| **Soil P (Mehlich 3) mg kg<sup>-1</sup>** | 22.6 | 175.0 | 46.3 | 348.6 |
| **Soil K (Mehlich 3) mg kg<sup>-1</sup>** | 83.1 | 265.0 | 71.5 | 199.8 |
| **Soil Al (Mehlich 3) mg kg<sup>-1</sup>** | 1579.7 | 1570.0 | 2838.8 | 1216.2 |
| **ISP<sub>1</sub> (environmental index %)** | 1.4 | 11.1 | 1.6 | 28.7 |
| **Texture** | Sandy loam | Fine sand | Sandy loam | Loam |
| **Minimum dose (kg ha<sup>-1</sup>)** | 0.00 | 0.00 | 0.00 | 0.00 |
| **Maximum dose (kg ha<sup>-1</sup>)** | 300.00 | 200.00 | 200.00 | 300.00 |

\* Trial used for economic optimal and agronomic optimal doses computation. N: nitrogen, P: phosphorus, K: potassium, Al: aluminium, ISP: phosphorus saturation index

Because soil grain-size distribution and organic matter content are compositional data, they were transformed into isometric log-ratios (ilr) to avoid self-redundancy, non-normal distribution and scale dependency [69]. The ilr transformation consists of log ratios of the geometric means of hierarchically-arranged components and groups of components, and can be interpreted as balances [70]. The hierarchical arrangement of components follows a balance scheme where balances split groups of components sequentially until each group contains a single part. Each balance is computed as in equation 5:

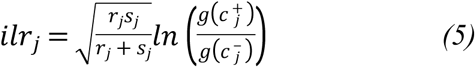

where for the j^th^ balance in [1, …, D-1] (D is the length of the compositional vector), r_j_ is the number of parts on the left-hand side, s_j_ is the number of parts on the right-hand side, c_j_^-^ is the compositional vector at the left-hand side, c_j_^+^ is the compositional vector at the right-hand side, and g() is the geometric mean function.

The proportion of the textural components and the carbon content formed the soil texture simplex. The balances are presented in Table 5. We followed the [denominator parts | numerator parts] notation [71].

**Table 5.**
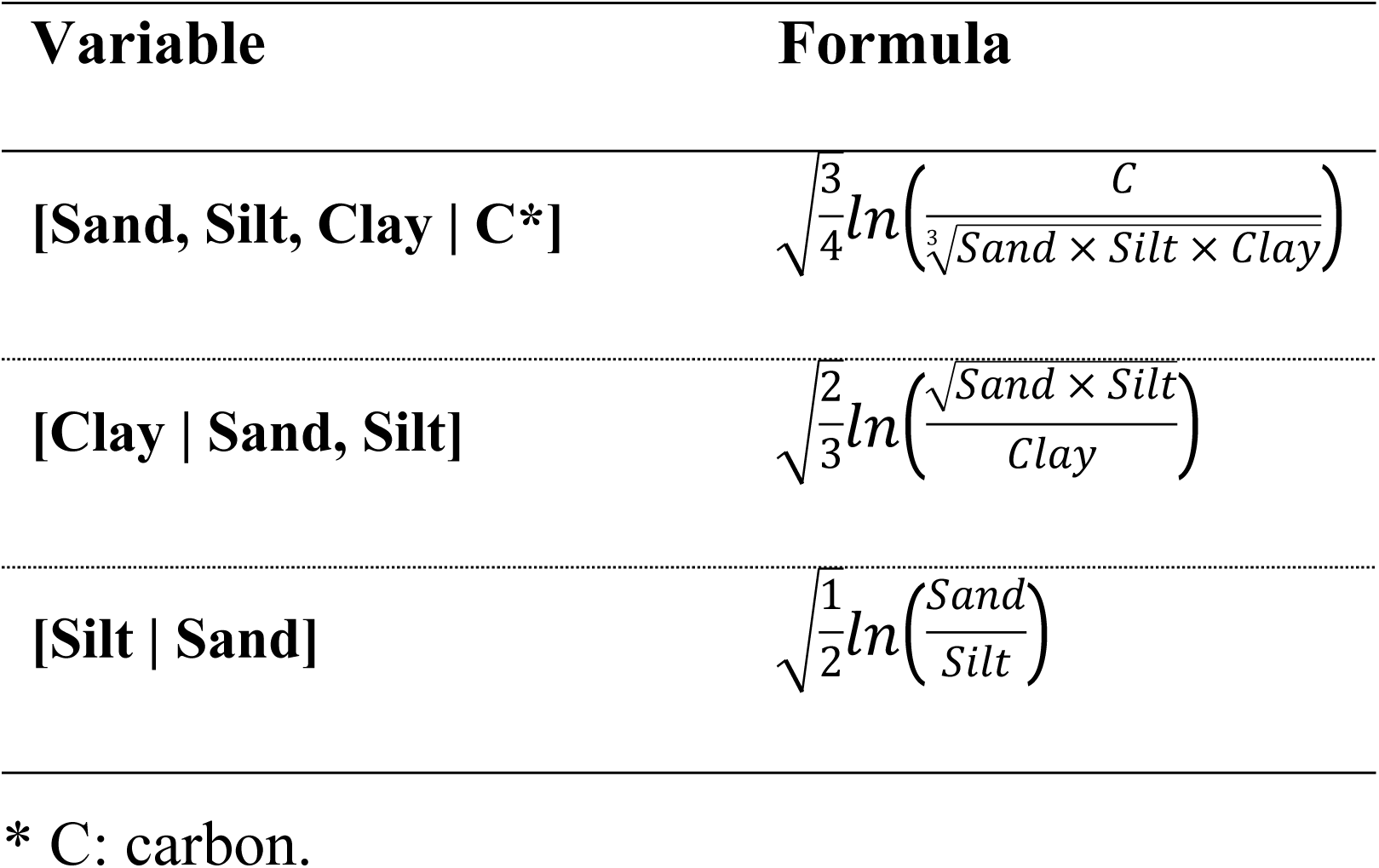
Soil texture variables used in the models.

Soil chemical compositions were partitioned into two simplexes (P, Al and K, Ca, Mg). The ilr variables were [Fv | Al, P], [Al | P] on the one hand and [Fv, Mg, Ca | K], [Fv | Mg, Ca], [Mg | Ca] on the other. Continuous expressions for soil types as used by Parent et al. [7] *i.e.*, poorly-drained loam, poorly-drained sand and well-drained sand, were balanced as [Gleyed | Podzolized] and [Loamy gleyed | Sandy gleyed] [56].

### 3.3 Weather data

Weather data were collected from the Environment Canada information system [72] using geographical coordinates for each site. The selected weather indexes were the cumulative precipitation – PPT, the Shannon Diversity Index for rainfall distribution – SDI [73] and the number of growing degree days – GDD using 5°C as baseline temperature. Weather variables were computed as in Parent et al. [7] (Table 6). We computed weather indices for the period between planting and harvest dates using the historical weather data for the past 5 years at each site.

**Table 6.**
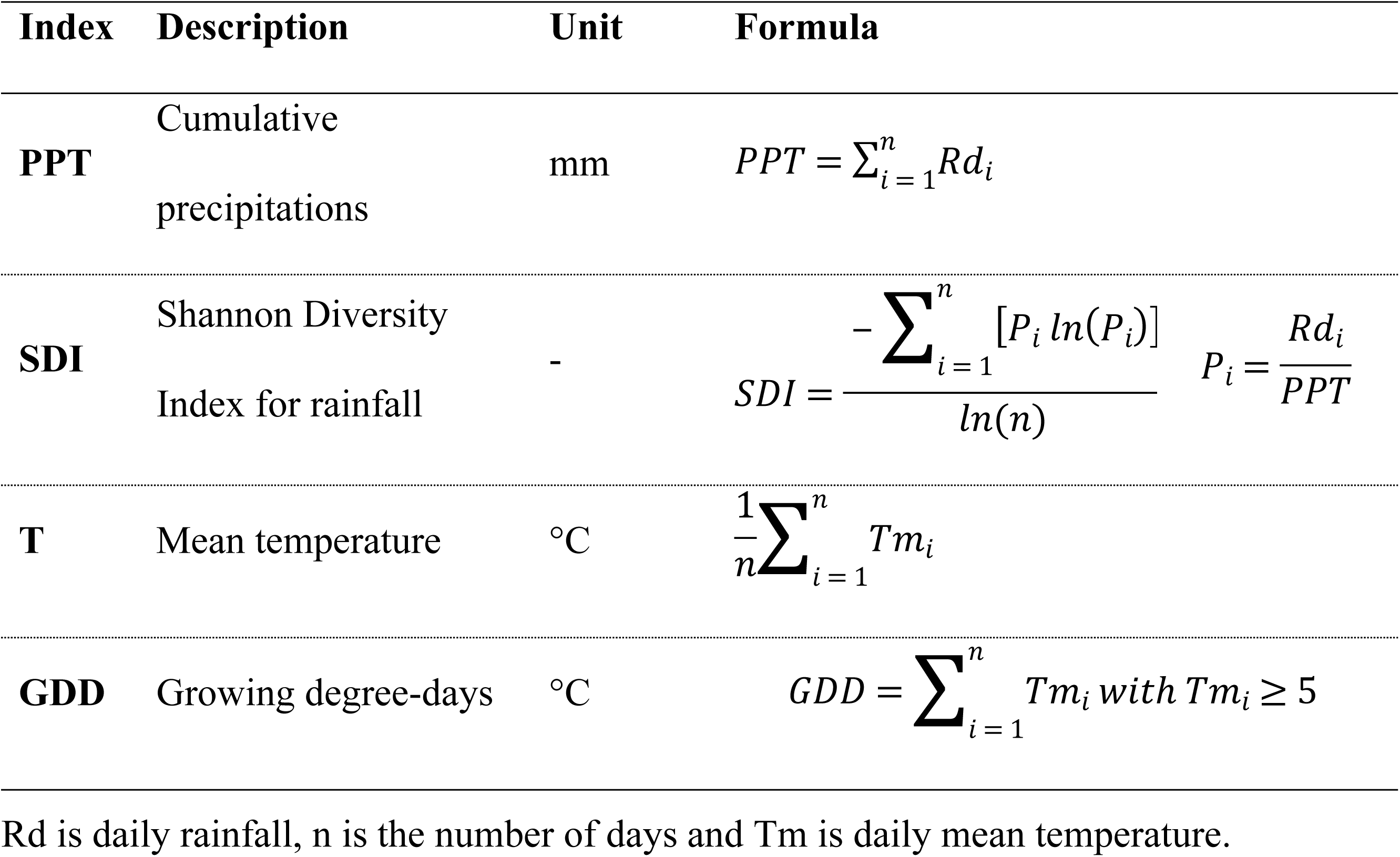
Equations to compute climatic indices.

| Index | Description | Unit | Formula |
| --- | --- | --- | --- |
| **PPT** | Cumulative precipitations | mm | $PPT = \sum_{i=1}^n Rd_i$ |
| **SDI** | Shannon Diversity Index for rainfall | - | $SDI = \frac{-\sum_{i=1}^n [P_i \ln(P_i)]}{\ln(n)} \quad P_i = \frac{Rd_i}{PPT}$ |
| **T** | Mean temperature | °C | $\frac{1}{n} \sum_{i=1}^n Tm_i$ |
| **GDD** | Growing degree-days | °C | $GDD = \sum_{i=1}^n Tm_i \text{ with } Tm_i \geq 5$ |

Rd is daily rainfall, n is the number of days and Tm is daily mean temperature.

### 3.4 Selection of features

#### 3.4.1 Predictive features

The study focused on potato yield-impacting factors reported by Parent et al. [7]. Candidate variables were soil Mehlich-3 P, K, Mg, Ca, Al and Fe composition, soil pH, and soil profile classes expressed as balances across soil textural gradients and across gleization-podzolization processes as in Leblanc et al. [56]. The length of the growing season, the preceding crop categories, seeding density and N, P and K fertilizer dosages were used as land management variables. The average 5-yr temperature (T), PTT, GDD and SDI were used as weather features. We selected features that contributed most to predicting output variables using the python scikit-learn *ExtraTreesRegressor* algorithm [74]. The trials with maximum yield less than 28 Mg ha^-1^ were discarded to avoid extreme cases of diseases, management failures or catastrophic weather events.

#### 3.4.2 Target variables

The data set is a collection of several experiments with specific objectives. Target variables were total yield, yield fractions, and SG. We separated marketable yield fractions with respect to tuber size as follows: large (L), medium (M) or small (S) size. Because these three fractions must add up to 100% of the marketable yield, they were treated as compositions. These compositional variables were transformed into isometric log-ratios of large-size tubers divided by the geometric mean of small- and medium-size tubers [M, S | L], and medium-size tubers divided by small-size tubers [S | M]. Since analysis of compositional data based on log-ratios of parts is not suitable when zeros are present in a data set [75], we proceeded by firstly imputing zero observations [76], reported mostly for large-size tubers. The detection limit was fixed at 65%.

### 3.5 Data preprocessing

The data were partially preprocessed in the R 3.6.2 statistical computing environment [77]. The tidyverse 1.3.0 package [78] was used for general data handling and visualization. The compositions 1.40-3 package [79] functions helped to transform compositional data into isometric log-ratios, and the robCompositions 2.2.0 package [80] helped to robustly impute missing values. The replacement of zeros in tuber sizes was performed using zCompositions 1.3.3-1 package [76].

The data preprocessing continued in Python 3.8.1 software [81]. The data set used to model tuber SG was cleaned of outliers using the Python SciPy package version 1.4.1 [82]. We used a z-score *i.e*., a signed number of standard deviations by which the value of an observation or data point is above the mean value of what is being measured on the multivariate data set. The threshold of the score value was set at 3. The data were handled in Python using NumPy version 1.17.5 [83] and pandas 1.0.0 [84] libraries. The matplotlib 3.1.3 package [85] was used for data visualization.

All the quantitative variables were scaled and centered to obtain zero mean and unit variance. The categorical variables were encoded by declining their factors in binary columns, each of which was denoted by 1 to specify the membership of the group of the column, and 0 otherwise.

### 3.6 Training and testing data sets

Schemes for partitioning data into training and testing sets vary between studies. Fortin et al. [6] used 60% for training and 40% for testing. Parizeau [86] suggested 50%, 20% and 30% for training, validation and testing, respectively. Crisci et al. [87] used a 75%–25% split while Chantre et al. [88] used a 82%–18% partition for training and testing, respectively. In this paper, the corresponding total input/output data pairs were divided into 70% for training and 30% for testing and model accuracy assessment. Soman and Bobbie [89] found shorter learning times and highest accuracies with such split proportions.

Moreover, self-contained and representative data collection is an important step to ensure the sufficiency and integrity of the training data [90]. Thereby, we partitioned the data set according to whether the tested element was N, P K, factorial design or another element (Mg, Ca). Thereafter, data were split at block level to avoid testing models on blocks comprising training samples.

### 3.7 Training models

#### 3.7.1 Machine learning algorithms

Four machine learning models were trained to derive an optimal model: *k*-nearest neighbors (KNN), random forest (RF), neuronal networks (NN) and Gaussian processes (GP). Model parameters were tuned using the random search with cross-validation method (*RandomSearchCV*) of the scikit-learn library version v0.22.1 [74].

#### 3.7.2 Mitscherlich model

We used a Mitscherlich-related 3D response surface for three variables inspired by Dodds et al. [91] in the multilevel modeling scheme of Parent et al. [7]. The Mitscherlich-related multilevel response surface was used as a predictive model for comparison with machine learning algorithms. The model was trained using the following equation:

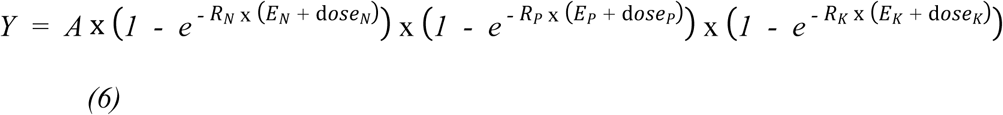

where *Y* is the target variable *i.e.,* marketable yield, *A* (for *Asymptote*) is the value of the target variable toward which the curve converges at increasing dosage, *E* (for *Environment*) describes the fertilizer-equivalent N (*E_N_*), P (*E_P_*) and K (*E_K_*) doses from the environment, and *R (Rate)* is the steepness of the curve relating each fertilizer equivalent environmental supply to *Asymptote*. The first-level parameters (*A*, *E* and *R*) were modeled as linear combinations of the predictors with random effect added to the intercept of the *Asymptote*. To make comparison with preceding models, the model performances were computed without any random effect (*level = 0*). The Mitscherlich multilevel model was fitted in R 3.6.2.

### 3.8 Evaluation of model performance

In all cases, the goodness-of-fit measure or predictive capacity of the developed models was based on the coefficient of determination (R^2^), the mean absolute error (MAE) and the root-mean-square error (RMSE). The R^2^ evaluates the proportion of variance in the target variable explained by the model as in equation 7:

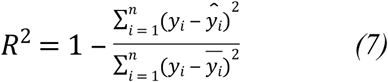

where *y*_i_ is the observed target variable value, 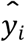 is the predicted target variable value, and 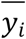 is the mean of observed target variable. The best possible score of R^2^ is 1 (or 100%), but the score may also be negative when the model is arbitrarily worse. Higher R^2^ values indicate less error variance. A constant model that always predicts the expected value of y disregarding the input features would yield a R^2^ score of 0 [74]. Typically, values greater than 0.5 are considered acceptable [92]. The MAE is the average of the absolute differences between predictions and observations as in equation 8:

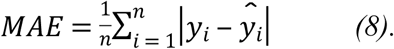

The MAE attributes equal weight to individual errors and is less sensitive than R^2^ or RMSE to large prediction errors. The RMSE is the square root of the average of squared differences between predictions and observations computed as in equation 9:

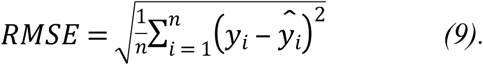

The RMSE attributes high weight to large errors due to squaring. Both MAE and RMSE indicate prediction errors in the units of variable of interest. Zero values indicate a perfect fit. Values less than half of the standard deviation of measured data were considered low [93]. The trained models were used to predict optimal N, P and K doses using some left-out experimental sites data.

### 3.9 Economic or agronomic optimal doses

The optimal nutrient input is the one returning yield of high-quality tubers [32], where profitability is maximized and the environmental footprint minimized [94, 95]. To compute the optimal economic N, P, K doses at a given site, all the predictive features, but not N, P and K doses, were held constant (fixed input data). The row of fixed input variables is stacked (reproduced) 1000 times to obtain a table with 1000 identical rows. We generated 1000 random N-P-K combinations of doses from uniform distributions of plausible doses varying between zero and 250 kg ha^-1^ for N, 110 kg ha^-1^ for P, 208 kg ha^-1^ for K. The table was altered in such a way that only N-P-K dosage changed following the random combinations.

A fertilizer cost was computed for each N-P-K triplet. Unit fertilizer costs were set at $1.20 CDN kg^-1^ for N, $1.10 CDN kg^-1^ for P and $0.90 CDN kg^-1^ for K. Tuber price was set at $250 CDN Mg^-1^ (1 Mg = 1000 kg). The difference between fertilizer cost and tuber revenue provided the marginal benefit from fertilizing. Economic optimal N-P-K dosage was reached where the net return was maximum. For tuber size and SG, an agronomic optimal N-P-K fertilizer dosage was deducted where the target variable reached a maximum.

No environmental footprint effect was used because of a lack of reliable sources, although they could have been implemented as an increase in the cost of unit dosage.

Our results are reproducible by using the codes, data and package requirements provided in a GitHub repository at https://git.io/JvYxd.

## 4 Results

### 4.1 Feature importance

The N fertilizer dose was by far the most informative feature in the marketable yield prediction models, followed by soil type, air temperature, length of growing season and soil texture. To predict large-size tuber yield ([M, S | L] balance), the N dose remained the most informative feature, followed by soil type and texture. Seeding density exceeded other features for medium-size tubers ([S | M] balance), followed by N dose, soil elements (P and Al Mehlich-3) and soil type. For tuber SG, weather indices, *i.e.*, Shannon diversity index, total rainfall and temperature, returned the highest scores (Fig 1). Preceding crops were not informative across target variables and were deleted before modeling.

**Fig 1:**
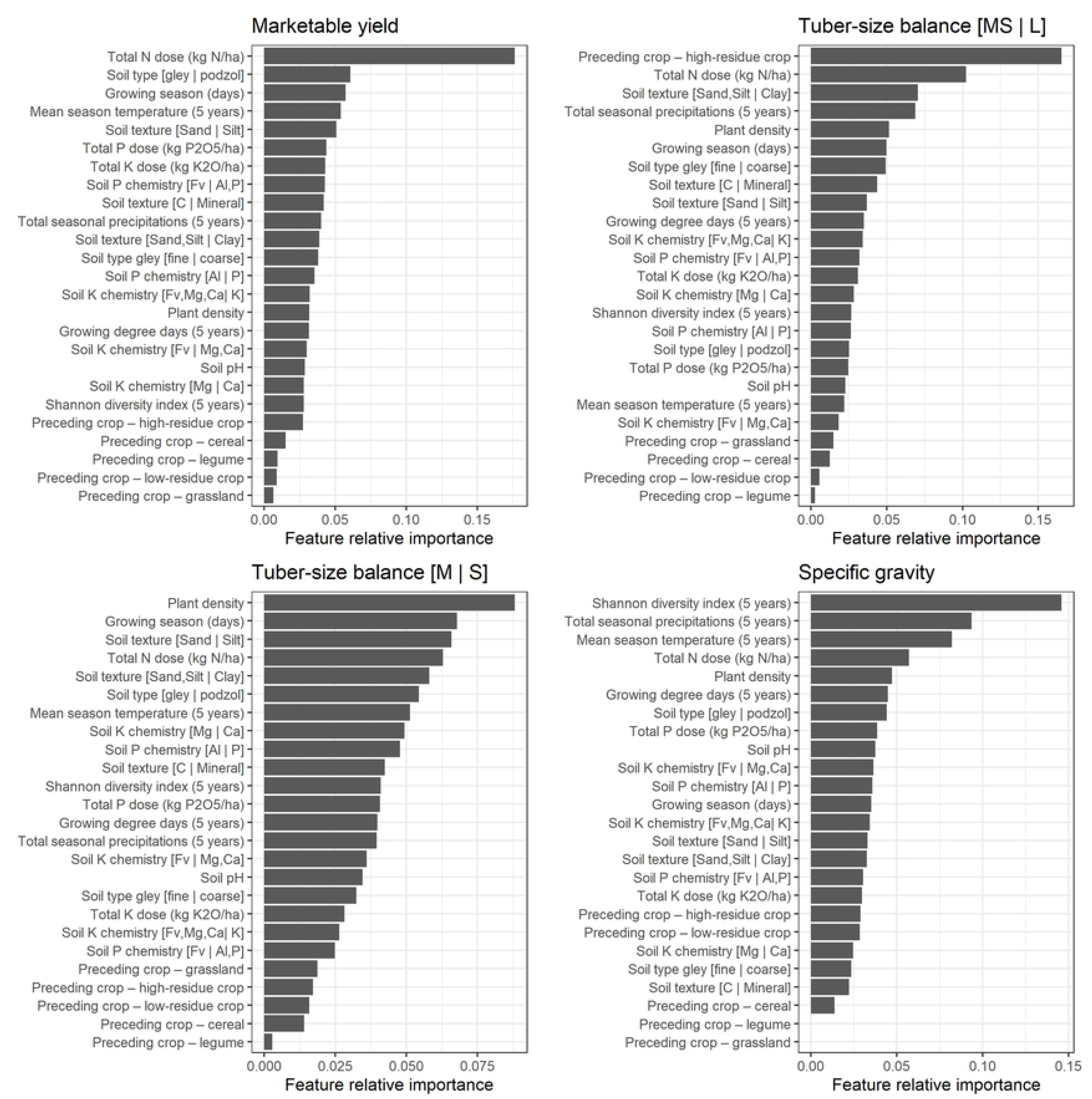
Predictive features importance for modeling.

The tuning parameters varied within the models depending on target variables. For the KNN models, the regressions were run with 19 nearest neighbors (k = 19) for yield, tuber size [M, S | L] balance and SG prediction models. For the [S | M] balance prediction model, k was set at 18 neighbors. The optimization procedure set the number of trees in the forest to 92, 12, 17 and 19 to train the RF models for yield, tuber size [M, S | L] and [S | M] balances, and SG, respectively. The number of features considered for splitting at each leaf node were selected automatically. The NN parameters were tuned to one hidden layer and a hyperbolic tangent activation function for all the target variables models. The tuned numbers of the hidden layer neurons were 100, 200, 100 and 200 for yield, tuber size [M, S | L] and [S | M] balances, and tuber SG respectively. The Matern covariance function was selected to train Gaussian processes for all the target variables and different noise levels: alpha was set at 0.195 for marketable yield prediction model, 0.136 for tuber size [M, S | L] balance, 0.031 for [S | M] balance, and 0.932 for tuber SG.

### 4.2 Comparison between models

Model performance to predict marketable yield, tuber-size balances and tuber SG was assessed using R^2^, MAE, RMSE, response curves shapes and economic optimal N-P-K dosage predictions for each model. For all the models, the predictive accuracy level was not affected after discarding the preceding crop classes.

#### 4.2.1 Goodness of fit

The model scores at training and testing for the different target variables are presented in Fig 2. There was a large gap between training and testing scores. The difference was lower for the Mitscherlich model, which also showed the lowest coefficient of determination and the highest MAE and RMSE. Its R^2^ values were 0.35 and 0.37 at training and testing, respectively. The R^2^ values of machine learning algorithm-based models ranged between 0.78 (NN) and 0.92 (KNN) at training, and between 0.49 (NN) and 0.59 (RF) at testing in predicting total yield. With the large-size tuber yield balance [M, S | L], the R^2^ values ranged between 0.72 (KNN) and 0.87 (RF) at training, and between 0.55 (KNN) and 0.64 (GP). The medium-versus small-size tuber [S | M] balance and SG prediction models were the most informative, as shown by the highest R^2^ values at both training and testing. The R^2^ values ranged between 0.83 (NN) and 0.93 (KNN) at training and between 0.62 (RF) and 0.69 (KNN) at testing in predicting small-size tuber balance, while for SG, they ranged between 0.72 (KNN) and 0.94 (RF), then between 0.58 (KNN) and 0.67 (RF) at training and testing, respectively. In general, model MAE and RMSE were slightly higher when R^2^ values were low. The practically-similar magnitudes between RMSE and MAE meant that all the individual differences between predictions and observations had equal weight.

**Fig 2:**
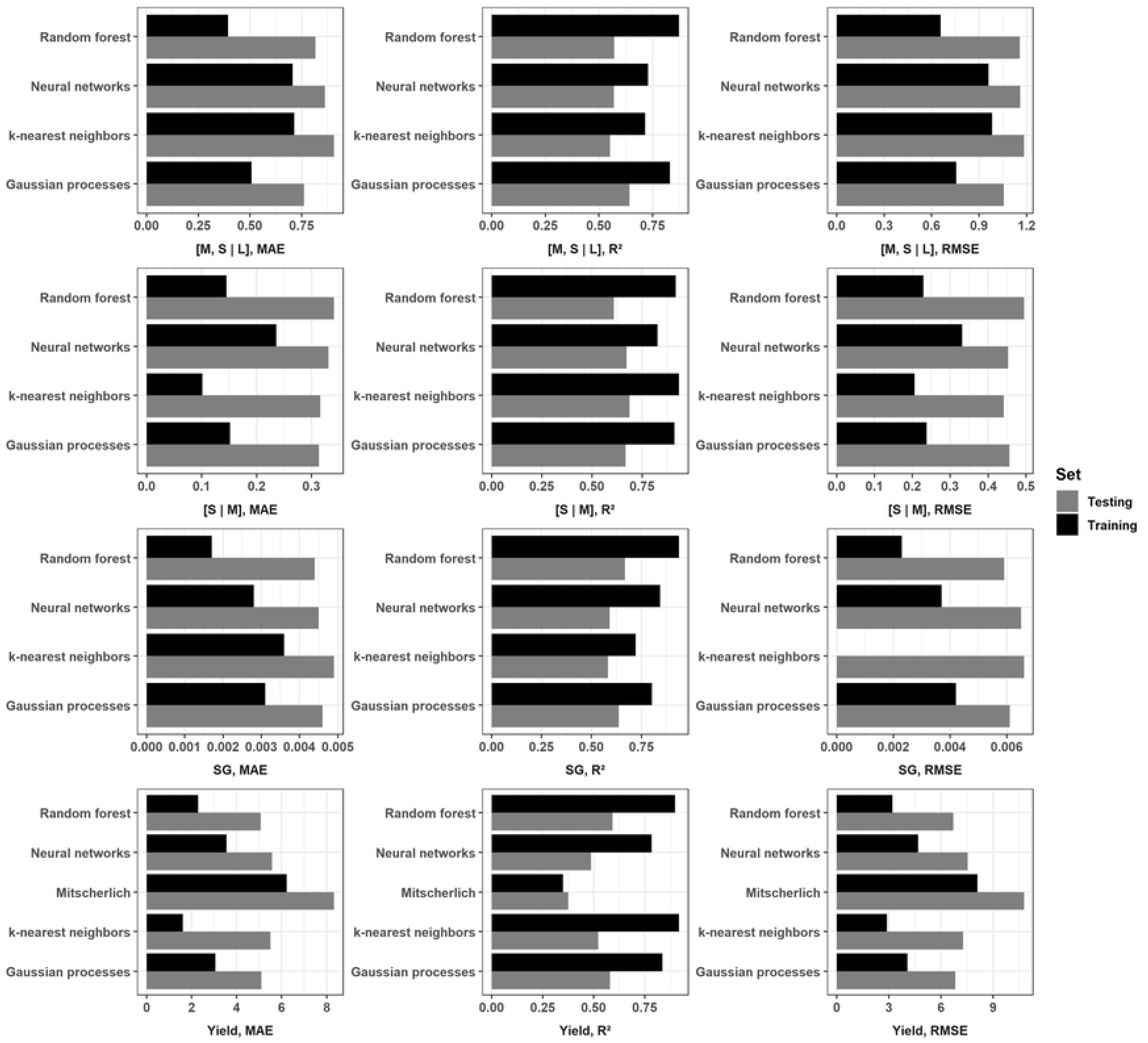
Comparison of models goodness of fit using R^2^, MAE and RMSE. R^2^: coefficient of determination, MAE: mean absolute error (Mg ha^-1^), RMSE: root mean square error (Mg ha^-1^), 1 Mg = 1000 kg. S: small, M: medium, and L: large size tuber

#### 4.2.2 Response curves

The marketable yield response curves are plotted in Fig 3 for each model with respect to the tested nutrient. There were disagreements between models. The Mitscherlich, NN and GP models generated smooth response curves, while the KNN and RF models generated stepped curves. The marketable yield was non-responsive to P application in the RF model. There was also no effect of K fertilization on the yield shown by the Mitscherlich and RF models. All models for the P trial somewhat underestimated marketable yield while response curves followed data for N.

**Fig 3:**
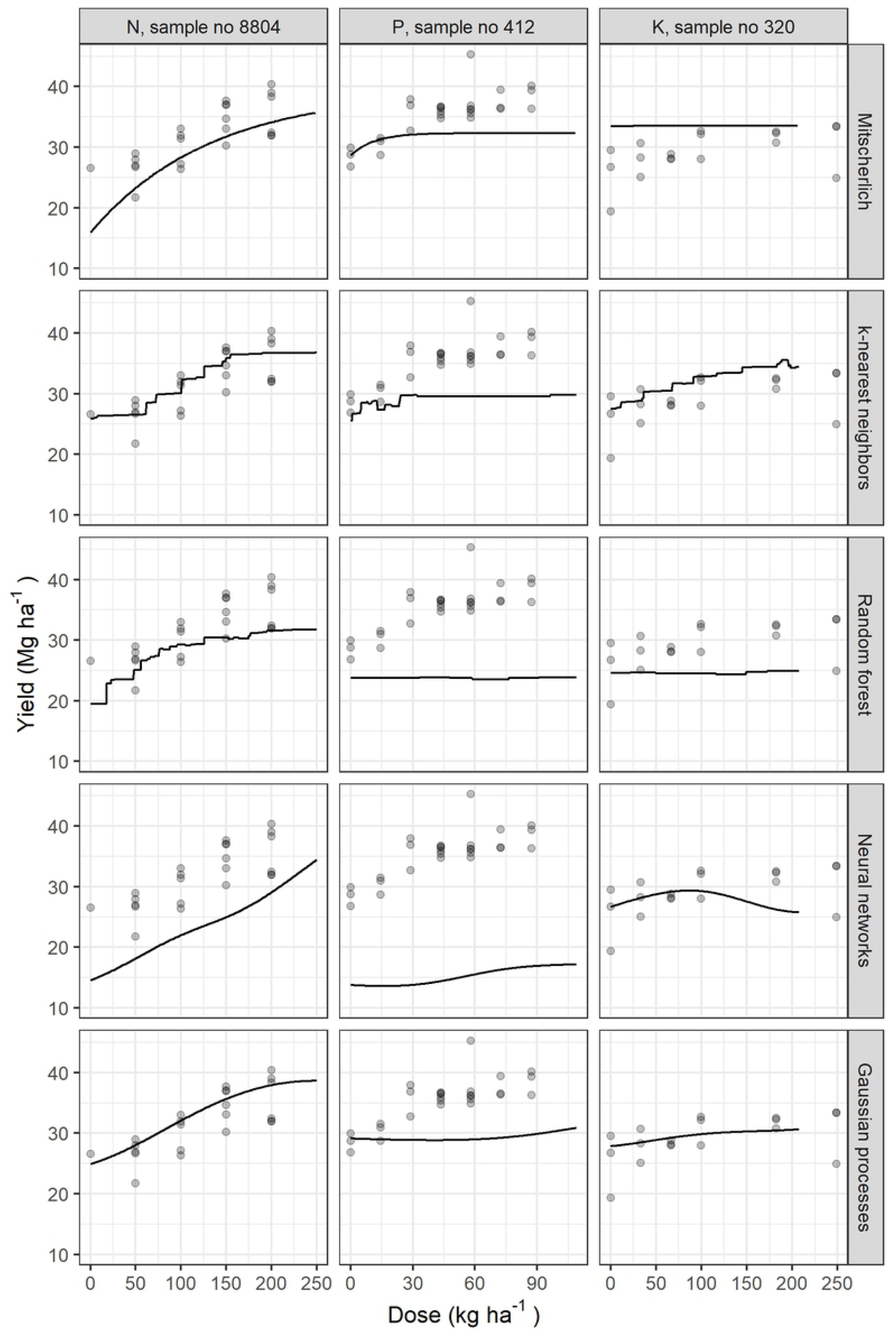
Examples of potato yield response to N, P or K fertilization using different models. N: nitrogen, P: phosphorous, and K: potassium

The Mitscherlich model was excluded for the analysis of other target variables. Figs 4–6 show how each model fits responses of tuber size balances ([M, S | L] and [S | M]), and SG, respectively, with respect to N, P or K dosage. The NN and GP models generated smooth curves, while the KNN and RF models generated stepped curves. The [M, S | L] balance (Fig 4) showed increasing response to N fertilization across models, while response was globally poor for P and K. For the [S | M] balance, responses increased with increasing fertilizer doses, except for P and K trials data fitted with GP model (Fig 5). There was also poor response for K trial with SG (Fig 6). The SG response decreased from zero K levels and increased then decreased as P dosage increased. For N trials, SG slightly increased then decreased as N dose increased in the RF model, but was non-responsive with the other models.

**Fig 4:**
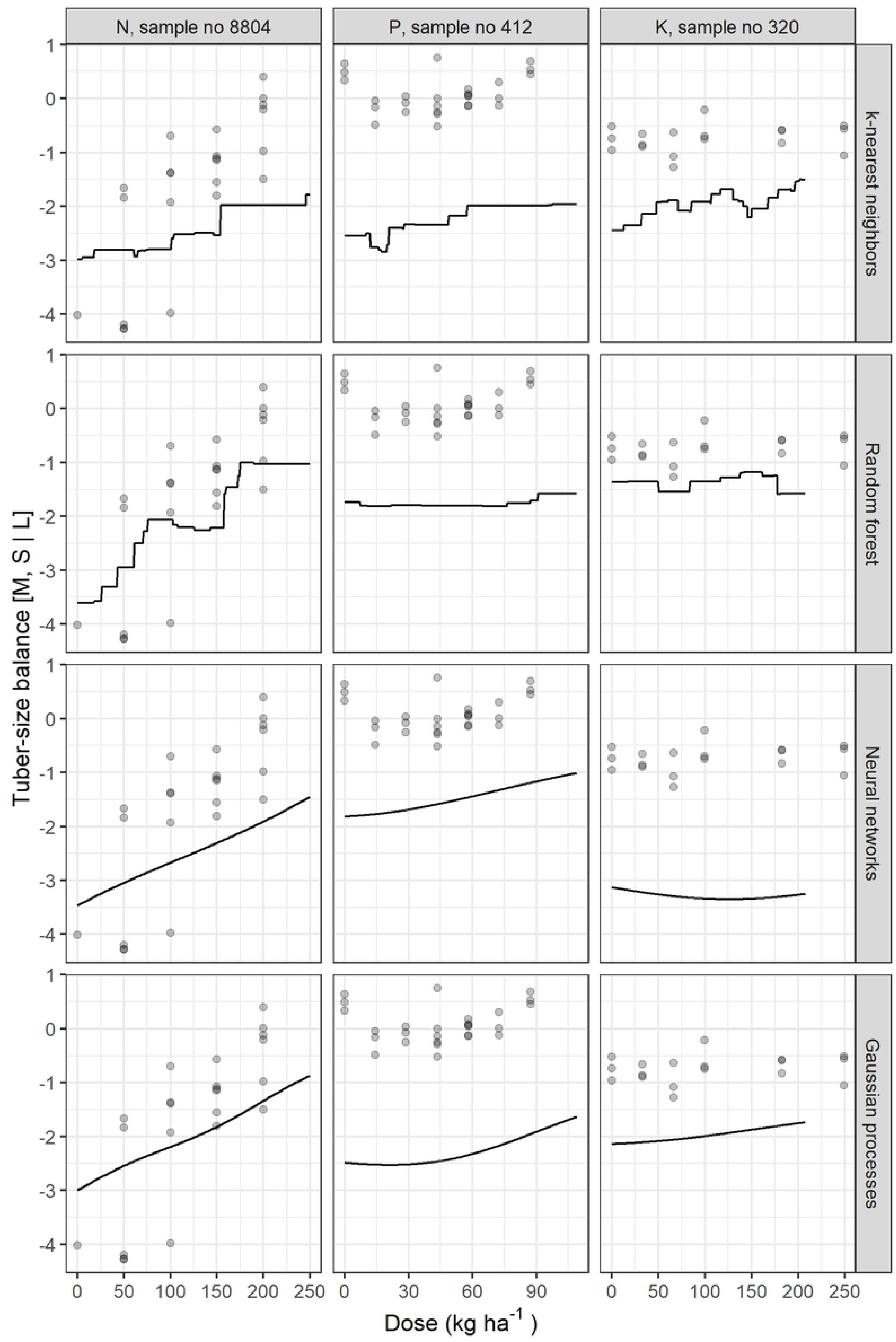
Examples of potato tuber size [M, S | L] balance response to N, P or K fertilization using different models. N: nitrogen, P: phosphorous, and K: potassium

**Fig 5:**
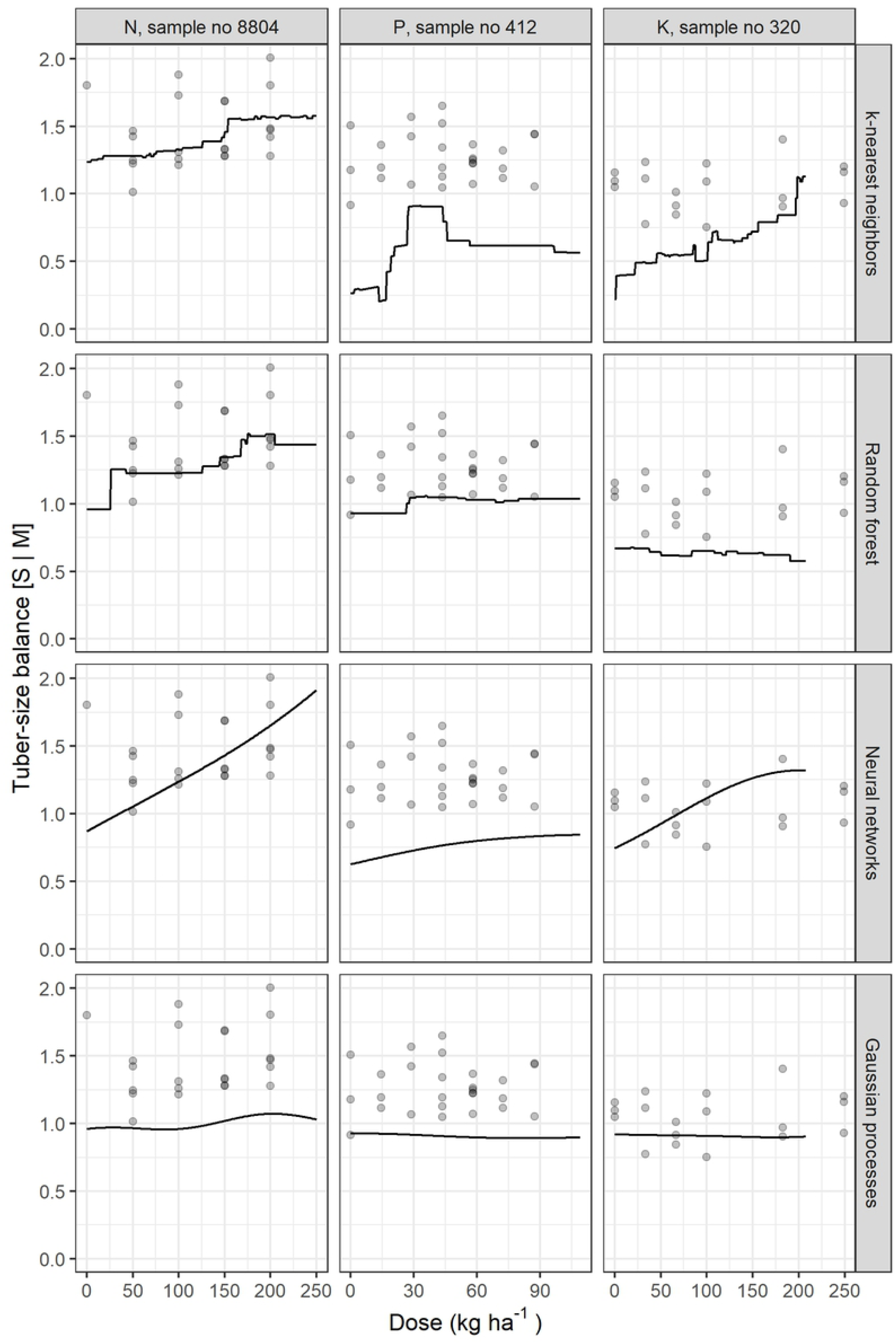
Examples of potato tuber size [S | M] balance response to N, P or K fertilization using different models. N: nitrogen, P: phosphorous, and K: potassium

**Fig 6:**
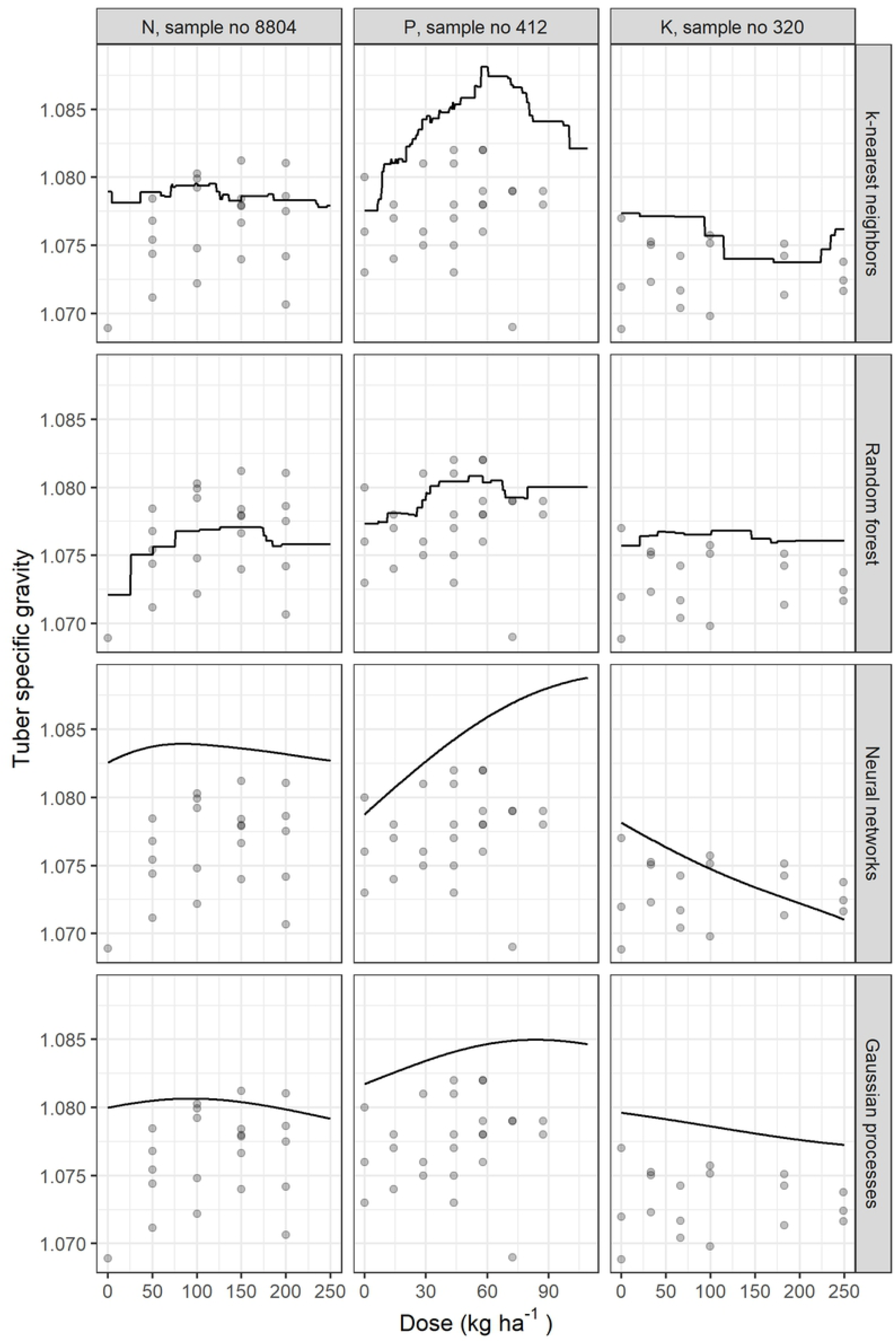
Examples of potato tuber SG response to N, P or K fertilization using different models. N: nitrogen, P: phosphorous, and K: potassium

#### 4.2.3 Predictions

The economic optimal doses and yield predictions and the agronomic optimal doses at maximum target values for tuber size balances and SG are presented in Fig 7.

**Fig 7:**
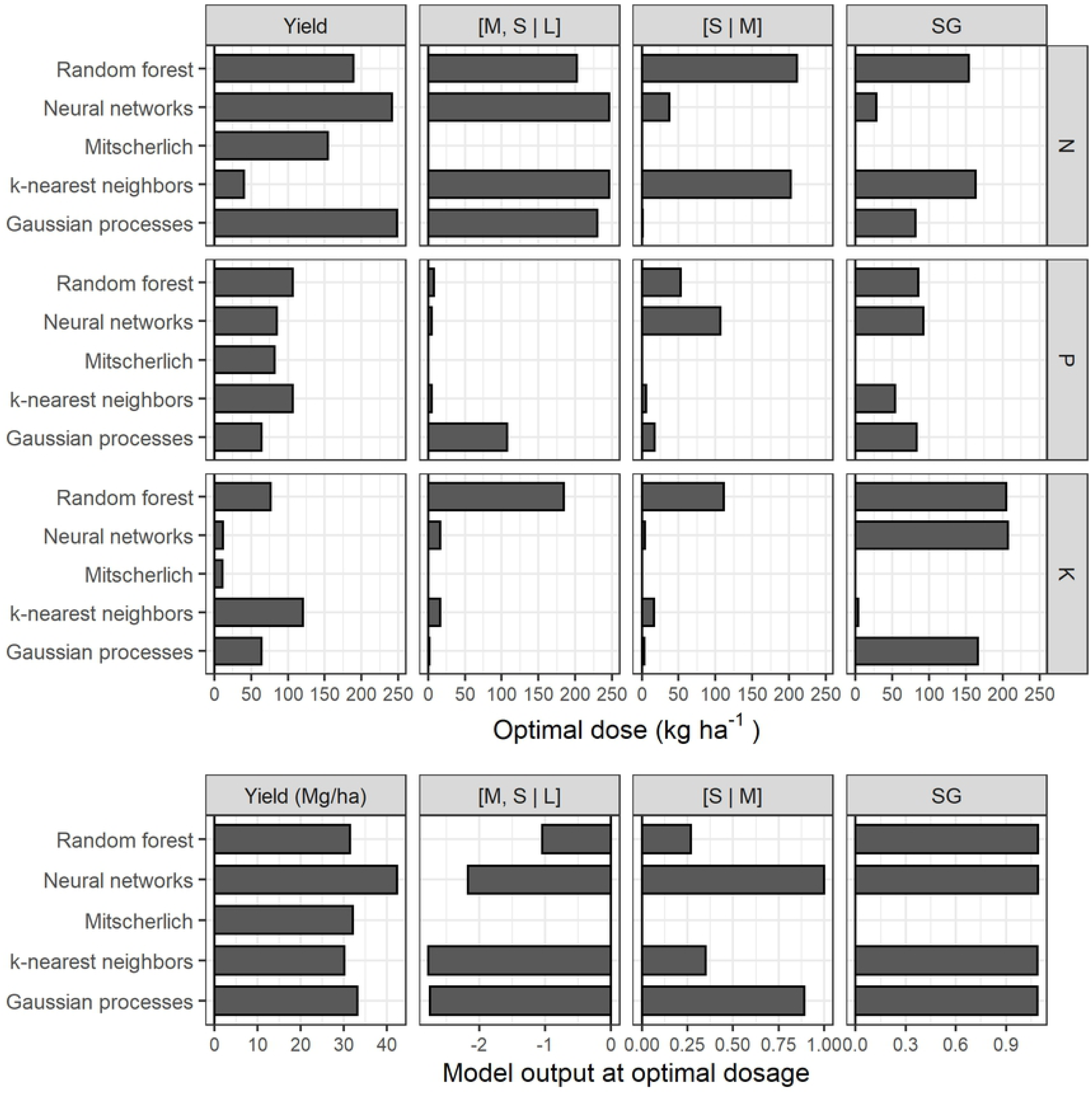
Economic optimal doses and yield predictions and agronomic optimal doses at maximum tuber size balance or SG for each model with a random selected test trial (N° 194). Fertilizer cost: N = $1.20 CDN kg^-1^, P = $1.10 CDN kg^-1^, K = $0.90 CDN kg^-1^ and crop price = $250 CDN Mg^-1^. Yield (Mg ha^-1^), N (nitrogen kg ha^-1^), P (phosphorus kg ha^-1^), K (potassium kg ha^-1^), [M, S | L] (medium and small versus large-size tubers log-ratio, unitless), [S | M] (small versus medium size tubers log-ratio, unitless)

The predictions varied with the model and the target. The Mitscherlich and NN models predicted negligible economic optimal K doses (11 and 12 kg ha^-1^ respectively) in marketable yield prediction models, while the site Mehlich-3 K level was classified as very low (83.1 mg kg^-1^) according to local standards [67]. The RF model suggested the highest cumulative agronomic optimum fertilizer doses, although its outputs were not the highest. With the tuber size [M, S | L] balance prediction model, practicable doses were recommended only by the GP model for P (107 kg ha^-1^) and the RF model for K (185 kg ha^-1^), a scheme that is almost similar to the [S | M] balance prediction models. For this output, the GP model recommended only 17 kg P ha^-1^, while N and K were impracticable (1 kg ha^-1^ and 4 kg ha^-1^, respectively). Despite the extremely low environmental risk for P and the low level of soil K, some models predicted negligible doses of P and K mainly for tuber size balances.

### 4.3 Probabilistic predictions

In addition to point estimates shown by each model, the GP model can return posterior samples. Each sample is a function from which we can compute an economic optimal (marketable yield) or agronomic optimal (size balances or SG) fertilizer dose. Figs 8–11 present the results of 1000 generated samples for each target variable for the selected N, P and K trials. The average GP curve is shown as a black line, with its optimal dosage as a black dot. Five sampled GP curves are plotted as grey lines, with their optimal doses as grey dots. The probability distributions of the 1000 optimal doses are shown under the respective response curves. The figures show that predicted means of optimal dosage (black dot) did not always correspond to the most likely dosage (highest histogram bar) computed after running the sampling process. With yield prediction models (Fig 8), the mean economic optimal dose corresponded to the probabilistic prediction only for the N trial (250 kg N ha^-1^). For the tuber size [M, S | L] balance (Fig 9), the probabilistic prediction was equal to the mean GP prediction for P trial *i.e.*, 87 kg P ha^-1^, while N and K trials returned equal predictions with the [S | M] balance prediction models with 0.0 kg ha^-1^ and 0.70 kg ha^-1^, respectively (Fig 10). For tuber SG prediction models, none of the probabilistic recommendation matched the mean GP optimal dosage (Fig 11).

**Fig 8:**
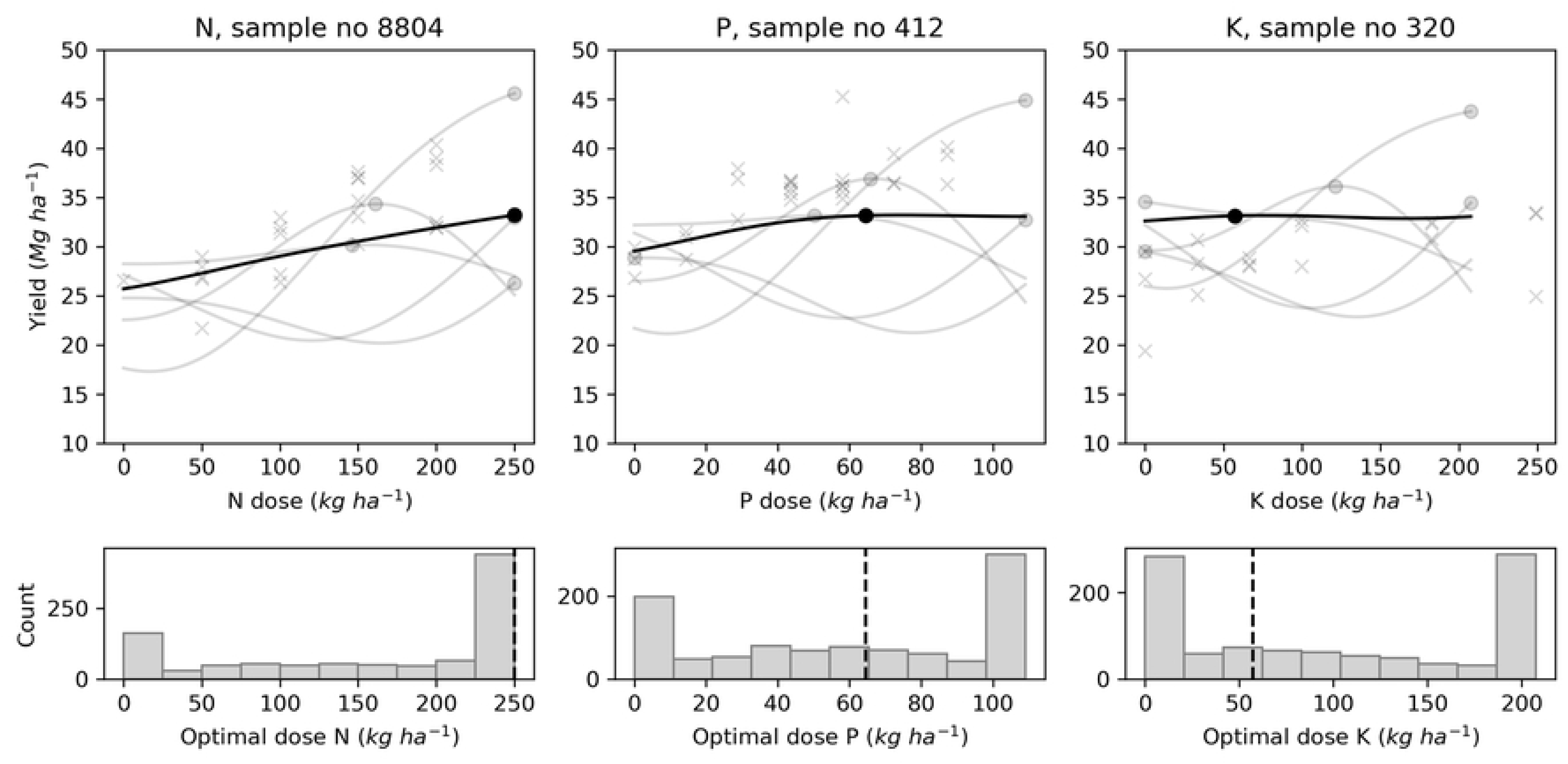
Examples of optimal economic N, P, K doses distribution with Gaussian processes using marketable yield for selected trials. N: nitrogen, P: phosphorous, and K: potassium

**Fig 9:**
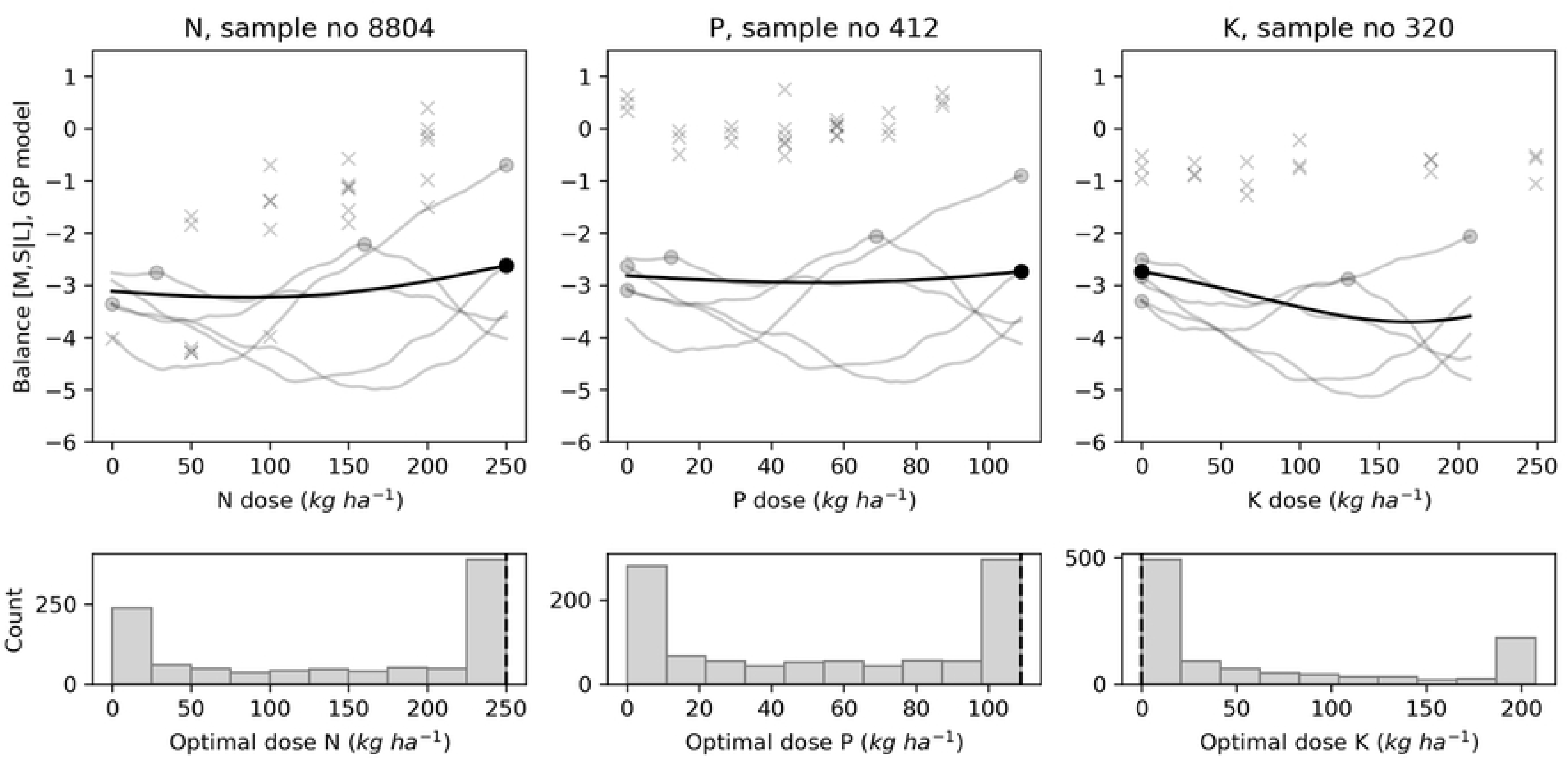
Examples of agronomic optimal N, P, K doses distribution with Gaussian processes using tuber size [M, S | L] balance for selected trials. N: nitrogen, P: phosphorous, and K: potassium

**Fig 10:**
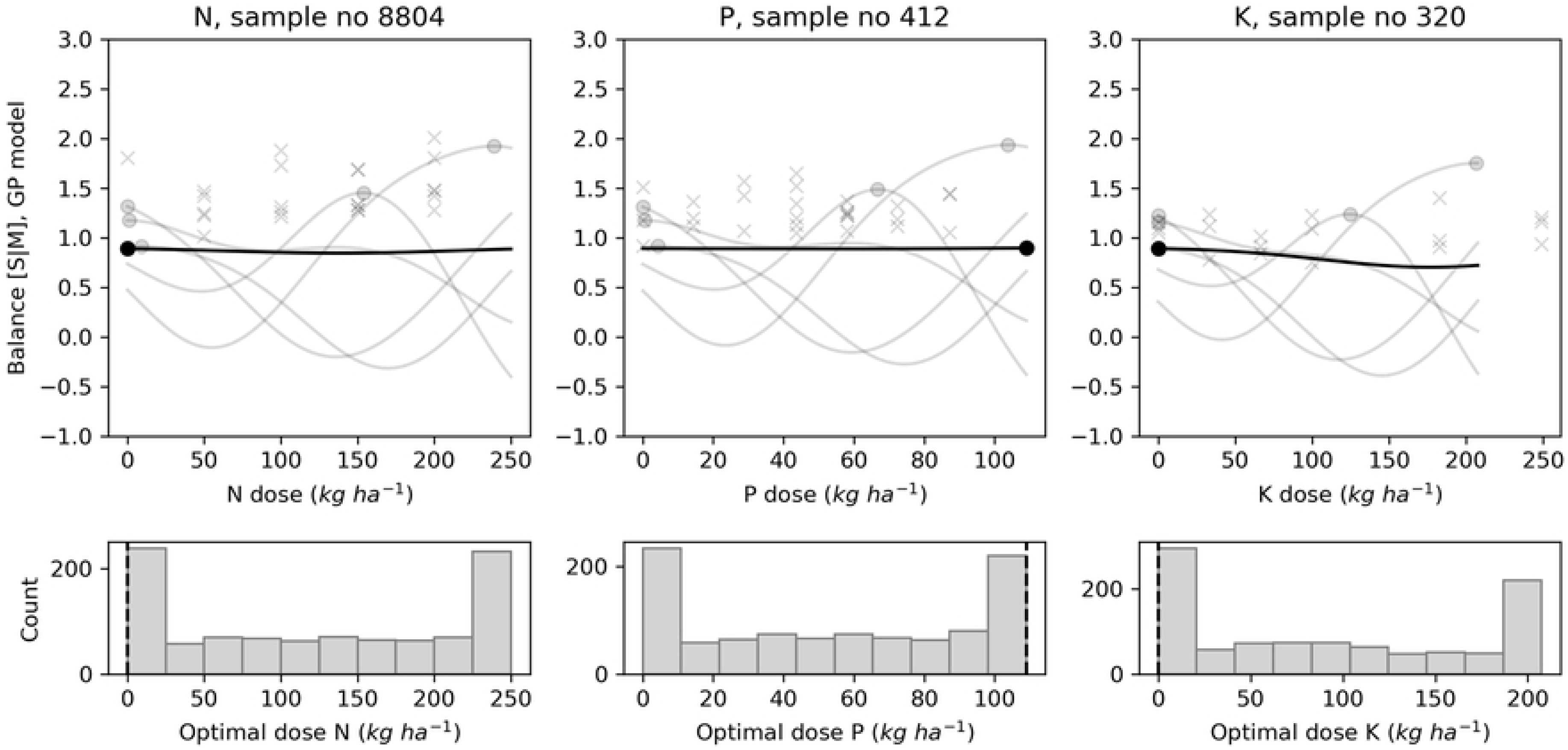
Examples of agronomic optimal N, P, K doses distribution with Gaussian processes using tuber size [S | M] balance for selected trials. N: nitrogen, P: phosphorous, and K: potassium

**Fig 11:**
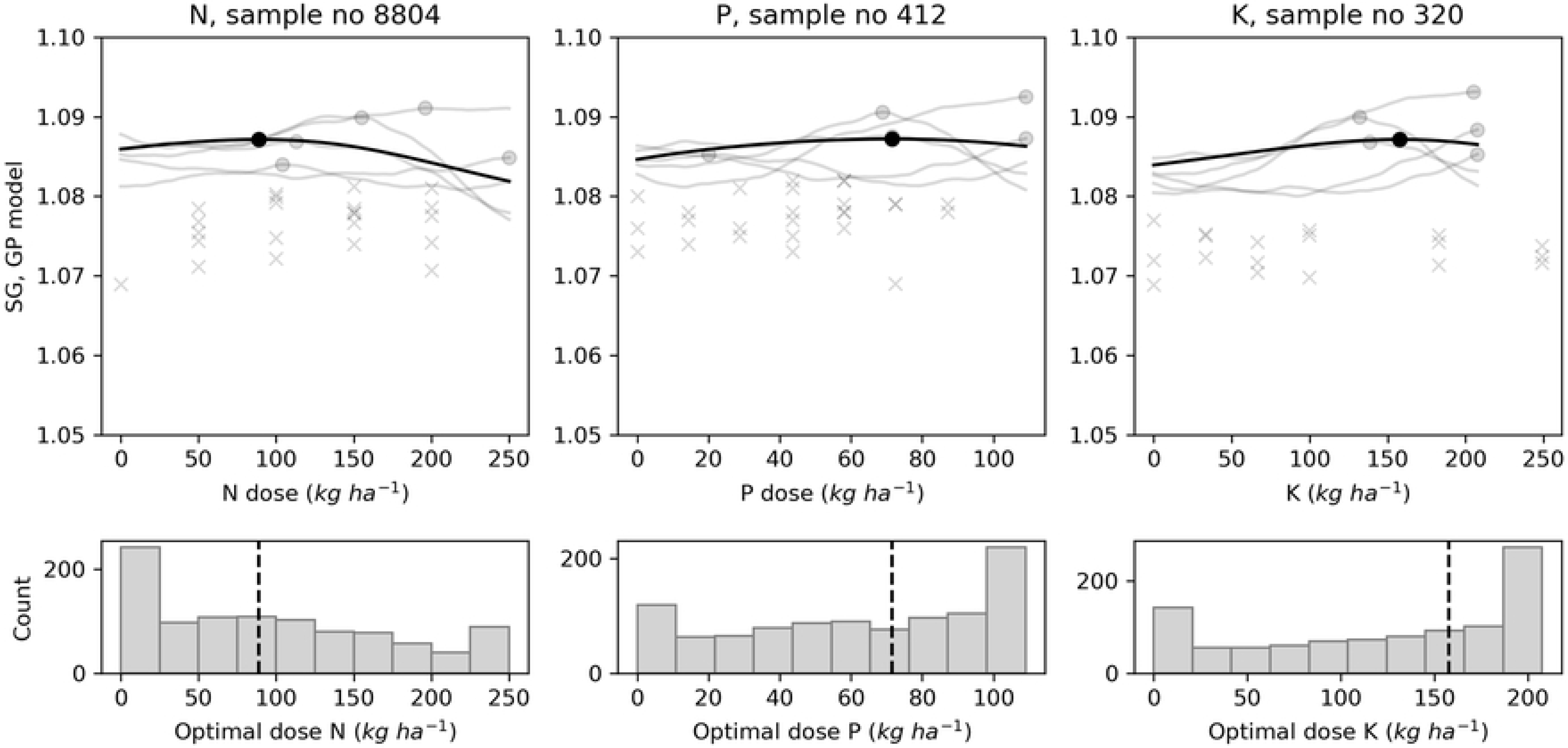
Examples of agronomic optimal N, P, K doses distribution with Gaussian processes using tuber SG for selected trials. N: nitrogen, P: phosphorous, and K: potassium

## 5 Discussion

### 5.1 Selection of features

The feature selection function selects a subset of variables for a learning algorithm to focus attention on the subset, especially when dealing with a large number of explanatory variables. The model-based approach incorporates the correlation structure between predictors and provides scores that indicate how useful or valuable each feature is in model building. Features with low or no importance could be removed without affecting model performance [74]. The preceding crops categories *i.e.*, grassland, small grains, legumes, low-residue crops and high-residue crops, as categorized by Parent et al. [7], returned zero (for tuber SG) or faintest scores (for other target variables) and were thus removed despite a substantial body of literature on the advantages of crop rotation to the next crop. Nonetheless, Zebarth et al. [96] stated that the amount of nitrogen mineralized from organic matter during the growing season cannot be predicted accurately. Torma et al. [97] found that the N supplied by soil and crop residues (maize, potato, silage maize, soybean, sunflower, winter rape, winter wheat) ranged from 20 to 132 kg ha^-1^, while the phosphorus ranged from 2 to 24 kg ha^-1^ and potassium from 13 to 218 kg ha^-1^. Rangarajan [98] stated that nutrient availability to the next crop depends on whether the entire plant or only the root system is left in the field, and on how environmental conditions govern the rate of organic matter decomposition.

For marketable yield and tuber size balances prediction models, the N dose was the most informative feature, probably because of its close relation to photosynthesis [99]. Applied in excess, it delays tuber maturity, stimulates foliage production, increases plant susceptibility to diseases and reduces tuber SG [100]. Crop yield is also determined by environmental conditions driving the physical, chemical and biological reactions [101] that are important in empirical or mechanistic models [4, 7–9, 102].

The selection process retained soil profile characteristics and weather events as major features. Levy and Veilleux [103] reported the effects of air and soil temperatures on potato growth mechanisms and tuber yield. Leblanc et al. [56] pointed out soil drainage conditions for loamy-gleyed profiles (poorly-drained loam), sandy-gleyed profiles (poorly-drained sand) and sandy-podzolized profiles (well-drained). Soil compaction has a negative impact on root extension and water movement *i.e.*, the reduction of nutrient uptake potential leading to a severe reduction of tuber yield [8]. Xu et al. [104] developed pedotransfer functions for potato grown on light-textured soils that could be useful in future models.

Dry matter production of potato crops is determined by the length of the growth cycle [105], which turned out to be a valuable feature. Camire et al. [106] stated that long growing season favors high-yielding late-season cultivars. Rex [107] found a close relationship between delayed harvest date and total yield, main-size marketable tubers and SG.

Seeding density was the most informative feature of the medium- to small-size tubers balance. Seeding density differentiates the number of tubers harvested, the weight of the tubers and the size distribution; higher plant densities promote higher yields in small and medium sizes [107–109].

The feature selection algorithm showed the impact of weather indices on tuber SG. The Shannon diversity index, total rainfall and temperature yielded the highest scores in a decreasing order. Al Soboh et al. [110] reviewed the factors affecting SG loss in crops of crisping potato and stressed that irrigation during early growth stages increases tuber dry matter content. Specific gravity could be reduced substantially if heavy rain occurred at the end of the season before harvest. They stated that potatoes grown during a period of increasing day length, temperature and light intensity produce tubers of high SG. In this study, GDD considered only daily mean temperatures higher than or equal to 5 °C as used by Parent et al. [7]. Moulin et al. [111] used a baseline of 7 °C and 30 °C as upper limit. Moreover, the general trend of SG response curves with respect to fertilization supported the results of Belanger et al. [112], Zebarth et al. [19] and Laboski and Kelling [113]. Excessive application doses of N and K along with high soil levels of either nutrient may reduce SG. Phosphorous application may increase tuber solids when soil test P levels are low. Specific gravity was not influenced by the relatively high levels of N and P used by Dubetz and Bole [114], while Maier et al. [115] found contrasted effects between trials.

Our analysis focused on the variation of N, P and K dosage while the other site-specific factors are kept constant. However, any predictive feature, especially biotic factors (*i.e.*, length of growing season, the preceding crop categories and seeding density), could also be optimized with respect to target variables.

### 5.2 Comparison of models

The performance of a predictive model is evaluated at testing or with unseen data set. The goodness of fit refers to how closely the model-predicted values match the true or observed values. Overfitting occurs where models perform well at training and badly at testing, while underfitting characterizes a model performing badly in both training and testing. Except for the Mitscherlich model, the model scores at testing showed discrepancies with training, reflecting problems of overfitting or underfitting. The differences between R^2^ values were highest for marketable yield prediction (Fig 2), reaching 0.40 with KNN. Based on those gaps, one could argue that our models did not generalize well from training to testing data. However, we used a robust approach by comparing different algorithms, tuning the hyperparameters and tuning the models using cross-validation. The R^2^ values at testing varied with respect to target variables, but were practically similar between models. The models estimated the proportions of medium- and small-size tubers ([S | M] balance) more accurately than those of large-size tubers ([M, S | L] balance), probably because of the high number of zero weight values among large-size tubers (21%) compared to tubers of small (0.06%) and medium (0.4%) size, at the early stage of our analysis. Imputing zeros to deal with measures where the large size was completely absent [75] improved the prediction quality of this fraction. Except for the Mitscherlich model in predicting yield, the R^2^ values at testing were greater than 0.50 and could be considered acceptable according to Moriasi et al. [92] for complex systems.

The Mitscherlich model returned a lower coefficient of determination in tuber yield prediction and was discarded for quality analysis (tuber size balances and SG). The KNN, RF, NN and GP algorithms more accurately approximated the unknown functions explaining tuber yield given the predictive features. However, it was difficult to select the best model since scores were practically similar. Cerrato and Blackmer [116] and several others [117–121] described similar ambiguities using classical statistical models.

Figs 3–6 indicated that the calibration and generalization procedures returned smooth response curves for the Mitscherlich, NN and GP models for all the target variables. Except for the low R^2^ value of the former, the NN and GP models appeared more suitable for making inferences.

The prediction of optimum fertilizer doses and optimum or maximum outputs showed some disagreements for the case presented (Fig 7). There should be a single economic optimal dose or agronomic optimal dose at each site each year. Some models were more consistent than others in deriving optimal doses depending on the target variable. At extremely low predicted N, P or K doses, it could be challenging to manage the fertilization program at low economic risk for producers, who generally consider that the cost of over-fertilization is low compared to the cost of under-fertilization [37, 38]. The probabilistic prediction capability of Gaussian processes may help to determine credible dosage.

### 5.3 Probabilistic predictions

Sampling from a Gaussian process looks like rolling a die, returning a different function each time. Figs 8–11 showed only five possible functions for each target variable. By sampling the process numerous times, we generated a distribution of economic or agronomic optimal fertilizer doses as those shown by the histograms of the figures. The distributions often show frequent optima at the edges to the NPK grid, *i.e.*, at dose of 0 or 250 kg ha^-1^. This phenomenon emerges from sampling continuously increasing or decreasing GP samples, which are more frequent when the sample is close to patterns in data where the response to fertilizer is flat. A zero-fertilizer recommendation could be interpreted as a soil sufficiently fertile to supply the crop, or a soil poorly responsive due to other constraints [122] such as pests and diseases [20, 123] or weed damage [124]. Nevertheless, we covered a wide range of factors that may impact potato crop growth and yield without falling into mechanistic modeling. Fertilizer doses more than 250 kg ha^-1^ may be excessive, since the maximum limits according to local standards are 175 kg ha^-1^ for N, 87 kg ha^-1^ for P and 199 kg ha^-1^ for K [67].

To face predictions falling at the edges, the optimal fertilizer dosage could be selected within a range of conditional expectation as processed by Khiari et al. [43] when defining P optimal dose for acid coarse-textured soils. The *x^th^* conditional expectation dose is the optimal dose that produces optimal yield *x%* of the time. For example, the 60^th^ percentile would be the sampled optimal dose that produces optimal yield 60% of the time for a given site. Khiari et al. [43] assessed the 50^th^ and 80^th^ percentiles. The mean (50%), the median or any other percentile dose could be computed to support decision-making. For example, the mean GP and the probability distribution processes returned the upper bound of the simulation dosage (*i.e.*, 250 kg N ha^-1^) as the economic optimal dose for the N trial with the marketable yield prediction model (Fig 8). The conditional expectation percentiles showed that a lower dose (*i.e.*, 223 kg N ha^-1^) could be recommended, producing optimal yield 55% of the time. At the 60^th^ percentile or more, the full dose *i.e.*, 250 kg N ha^-1^ must be applied.

## 6 Conclusion

This study assessed machine learning techniques as an alternative for potato fertilizer recommendations at local scale usually handled by statistical models or meta-analysis at regional scale. A large collection of field trial data provided information to fit machine learning models with specific traits of cultivars, soil properties, weather indexes, and N, P and K fertilizers dosage used as predictive features. Five models, Mitscherlich, KNN, RF, NN and GP, were evaluated against optimal economic N, P and K doses derived from yield, or against optimal agronomic N, P and K doses derived from tuber size and SG. The models trained using machine learning algorithms outperformed the Mitscherlich tri-variate response predictive model. The marketable yield prediction coefficient (R^2^) varied between 0.49 and 0.59, while the Mitscherlich model returned 0.37. The large-size tuber balance was predicted with a coefficient varying between 0.55 and 0.64. The R^2^ varied between 0.60 and 0.69 in predicting medium-size tuber balance, and between 0.58 and 0.67 for SG. The N, P and K optimal doses could be recommended with respect to marketable yield, tuber size or SG using the NN and GP models, which appeared to be the most suitable for making inferences. Response surfaces were obtained by marginalizing the models using N-P-K doses generated from uniform distributions under constant weather conditions, soil properties and land management factors. The GP model stood up by its probabilistic framework in risk estimation for potato fertilizer recommendation in Quebec conditions.

As large amounts of data are being assembled into observational data sets, machine learning models may surrogate statistical models in making fertilizer recommendations in the context of precision agriculture. To assess model performance under real-world situations, it was an effective strategy to combine historical weather data since accurate future weather data covering the growing season are unavailable. We also focused on using easily-available features collected from routine analyses as predictors instead of mechanistic processes models. Any biotic factor other than fertilizer, *e.g.*, length of growing season or planting density, could be optimized with our model. Improvement will require more data from many more diverse environments and management scenarios. With more experiment data, the training and testing division could be performed at trial level to improve the model predictive ability. Moreover, since the data for this analysis were collected from small research plots, validation at production-scale fields is needed for decision making.

## 8 Supporting Information

**S1 Table. Quebec potato data set used for modeling.** ‘Potato_df.csv’ file available in ‘data’ repository at https://git.io/JvYxd.

